# A web bench for analysis and prediction of oncological status from proteomics data of urine samples

**DOI:** 10.1101/315564

**Authors:** Sherry Bhalla, Kumardeep Chaudhary, Ankur Gautam, Suresh Sharma, Gajendra P. S. Raghava

## Abstract

Urine-based cancer biomarkers offer numerous advantages over the other biomarkers and play a crucial role in cancer management. In this study, an attempt has been made to develop proteomics-based prediction models to discriminate patients of oncological disorders related to urinary tract and healthy controls from their urine samples. The dataset used in this study was obtained from human urinary peptide database that contains urine proteomics data of 1525 oncological and 1503 healthy controls with the spectral intensity of 5605 peptides. First, we identified peptide spectra using various feature selection techniques, which display different intensity and occurrence in oncological samples and healthy controls. Based on selected 173 peptide-based biomarkers, we developed models for predicting oncological samples and achieved maximum accuracy of 91.94% with 0.84 MCC. Prediction models were also developed based on spectral intensities with known peptide sequences. We also quantitated the amount of protein in a sample based on intensities of its fragments/peptides and developed prediction models based on protein expression. It was observed that certain proteins and their peptides such as fragments of collagen protein are more abundant in oncological samples. Based on this study, we also developed a web bench, CancerUBM, for mining proteomics data, which is freely available at http://webs.iiitd.edu.in/raghava/cancerubm/.

## 1 Introduction

Cancer, one of the leading causes of mortality in the world, is responsible for millions of premature deaths. More than 11 million people die from cancer every year and with the present data, it has been estimated that 11.4 million people will be dying of cancer in 2030[1]. The mortality due to cancer can be reduced significantly, if cancer cases are detected and treated timely. Thus, diagnostics is as important as treatment of cancer[2]. This accounts for the recent research related to new and better cancer biomarkers from different sources of biofluids.

Biofluids are assumed to reflect the collection of tissues present within the human body[3]. Proteomics of body fluids (e.g., urine, plasma, and serum) can be used to discover novel biomarkers for early detection of cancer. Urine is one of the most assuring biofluids in clinical proteomics, as it is easy to collect, non-invasive and contains sufficient proteins and peptides[4, 5]. There are various techniques to analyze urine proteins and peptides such as two-dimensional electrophoresis with mass spectrometric and/or immunochemical identification of proteins (2DE/MS)[6-10], liquid chromatography coupled to mass spectrometry (LC/MS)[8, 11] and surface-enhanced laser desorption ionization mass spectrometry (SELDI/MS)[12] and capillary electrophoresis/mass spectrometry (CE/MS)[13]. 2DE/MS is applicable to large molecules, but not applicable to peptides, which are less than 10 kDa. CE/MS is most suitable technique for urine proteomics as it has a number of advantages over other techniques. In the past, attempts have been made to develop peptide-based cancer biomarkers from proteomics data of urine. The main limitation of previous studies is that they are based on small number of samples[5]. In order to develop reliable biomarkers, one should use large number of samples and proteomics data should be generated under identical conditions using the same technological platform[14].

Human urinary peptide database is a reproducible, high-resolution database for proteome analysis of naturally occurring human urinary peptides of 47 different pathophysiological conditions. This database includes 13,027 urine samples to date. The purposefulness of the database is to assist as a platform for the definition and validation of biomarkers for a variety of diseases and physiological changes[15]. The dataset from human urinary peptide database has been used previously for predicting biomarkers for chronic kidney diseases that resulted in 85.5% sensitivity and 100% specificity[14]. The subset of this database has been used to predict status of bladder cancer using panel of four peptides[16]. In another study, 12 peptide biomarkers for prostate cancer have been used to classify cancer patients and healthy individuals[17].

In the present study, we extracted proteomics data of around 3000 urine samples from human urinary peptide database. This proteomics data has been generated in identical conditions from a single platform. In this study, using spectral intensity of peptides as features, we identified biomarkers for oncological disorders on three levels, which are characterized by spectra (calibrated mass (Da) and CE time (min)), peptide (characterized by mass, CE time and peptide sequence) and protein quantity (mass, CE time, peptide sequence and protein). These biomarkers have been used for developing models for prediction of oncological status of patients from their urine proteomics data. Our main emphasis is to present major biomarkers associated with oncological disorders of urinary tract and to classify the oncological and healthy samples on the basis of these biomarkers derived from urine proteomics data. Finally, a web bench has been developed for providing service to the scientific community, particularly for those working in the field of cancer diagnostics. To the best of authors’ knowledge, there is no webserver where user can analyze their CE/MS data for identification of biomarker and prediction of oncological status.

## 2 Materials and methods

### 2.1 Data

We extracted CE/MS-derived intensity values of 1525 samples of oncological disorders, which include 526 patients of prostate cancer, 362 patients of benign prostatic hyperplasia, 334 patients of bladder oncological, 123 patients of kaposi sarcoma, 127 patients of renal cancer, 41 patients of prostatic intraepithelial neoplasia III and 12 patients of pheochromocytoma along with 1503 healthy controls from the 13,027 samples of the human urinary peptide database v3.0 (updated on February 11, 2011). The values in the database represent the calibrated amplitude of the mass spectrometric signal of 5605 peptides for oncological samples and healthy controls that are used for the further analysis and development of machine learning models.

### 2.2 Normalization of data

It was observed that the intensities of peptide spectra have wide range of variation, which is difficult to use for developing biomarkers. In order to mine this data, we performed normalization of data. We performed log transformation of intensities where we calculated natural log of intensities after adding value of 1 to the intensities.

### 2.3 Feature Selection

The selection of relevant features is one of the important tasks for developing any prediction model, as all features are not relevant. These unrelated features create noise or error in data mining. In this study, we have 5605 peptide spectra, thus we used the feature selection technique to reduce number of features. We used “cfsSubsetEval” attribute evaluator of popular Weka[18] package using “BestFirst” search method. This method resulted in 173 peptide spectra; we referred this feature set of 173 as fset173 (Figure 1).

**Figure 1:**
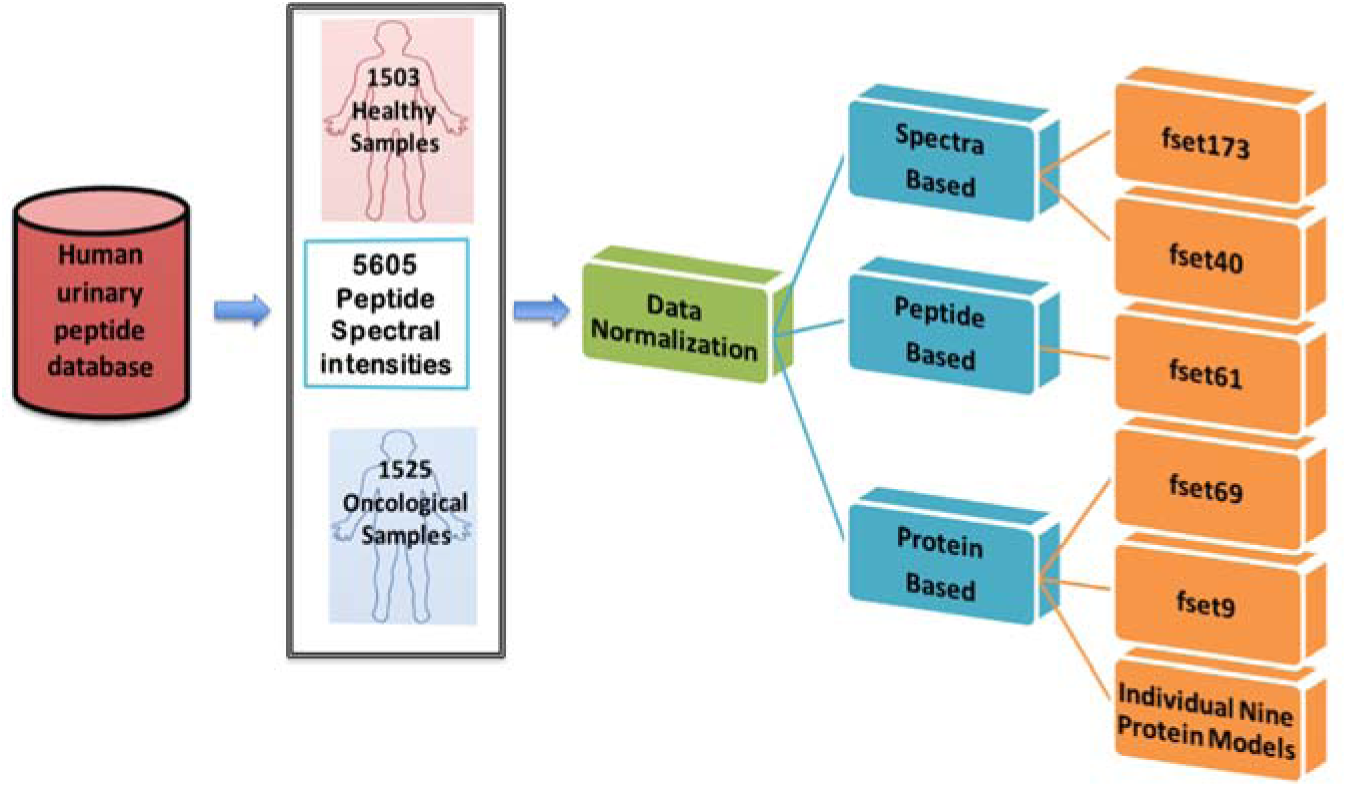
Peptide and protein feature sets created from human urinary peptide database for machine learning model development and analysis.

In addition to above feature selection technique, we also used a simple technique for feature selection. In this technique, we identified non-zero mass spectrometric signal for each peptide separately for oncological patients and healthy controls. Based on the difference in non-zero samples in oncological and healthy controls, we identified top 20 peptides that are most frequent in oncological patients than in healthy controls. Similarly, we identified top 20 peptide spectra that are more frequent in healthy controls than in oncological patients. This way we got a total of 40 peptide spectra used in this study for developing models. We referred to these 40 peptide spectra, selected based on frequency, as the fset40.

We extracted the protein information of the above spectra of peptides from human urinary peptide database for further analysis. Out of 173 peptides, information was available for 61 peptides with their amino acid sequence and their source proteins. We used intensity of these 61 peptides as features for developing models and referred to them as the feature set, fset61.

We also extracted the protein information of the 953 peptides from a total of 5605 spectra of peptides. We mapped these 953 peptides to human urinary peptide database, which eventually mapped to 118 proteins. There were 69 proteins out of 118 proteins having more than two peptide fragments. We called this as protein feature set and referred to as fset69.

We also mapped 61 peptides, in fset61, on proteins and obtained nine proteins having two or more peptides. We used these nine proteins for developing protein-based models and called these proteins as feature set, fset9. Nine individual models for each of the nine proteins were also built to ascertain the capability of each protein to differentiate between oncological and healthy samples.

**Protein Features**. We identified source protein of each peptide spectra; next challenge was to use these proteins for developing prediction models. Each protein in our dataset has two or more peptides and their spectral intensities. We computed expression of a protein based on intensities of peptides belonging to this protein. The intensity of a peptide is proportional to its abundance in sample or expression/amount of protein in the sample. Thus, we computed expression or amount of protein in a sample from spectral intensities of peptides of this protein, using mean, median and maximum values.

### 2.4 Machine Learning Techniques

In this study, most of these models have been implemented using popular software packages SVM*^light^* (http://svmlight.joachims.org/) and Weka. Weka contains different classifiers such as Random Forest[24] and SMO[25] *etc*. We applied these classifiers for the classification of oncological and healthy samples. We implemented SVM using SVM*^light^*.

## 3 Results

In order to mine vital oncological biomarkers from urine for various oncological disorders associated with the urinary tract, we have analyzed the human urinary peptide database. Based on observations, *in silico* models have been developed for the prediction of oncological status from proteomics data of urine samples. Following is the detailed description of models developed for discriminating oncological samples and healthy controls.

### 3.1 Prediction on the basis of peptide spectra

Our dataset contains 1525 and 1503 urine samples from oncological patients and healthy controls respectively; each sample has spectra of 5605 peptides. In order to select relevant peptides, software package Weka[18] has been used for feature selection, which identified 173 peptides spectra. These 173 peptides spectra are referred in this study as the feature set, fset173 (feature set of 173 peptides). The performance of models developed using log intensity of fset173 peptides has been shown in Table 1. As shown in Table 1, SVM-based models performed reasonably well, where Matthews correlation coefficient (MCC) of 0.84 was achieved.

**Table 1:** The performance of models developed based on different machine learning techniques using features in fset173.

| Method | Threshold | Sensitivity | Specificity | Accuracy | MCC |
| --- | --- | --- | --- | --- | --- |
| **Random-forest** | 0.5 | 92.59 | 87.03 | 89.83 | 0.80 |
| **SMO-Polykernel** | 0.1 | 88.79 | 88.76 | 88.77 | 0.78 |
| **SMO-Puk** | 0.1 | 95.02 | 79.51 | 87.32 | 0.76 |
| **SMO-RBF** | 1 | 99.8 | 19.16 | 59.78 | 0.32 |
| **SVM** | 0 | 93.18 | 90.69 | 91.94 | 0.84 |

**MCC: Matthews Correlation Coefficient**

In order to discriminate between urine sample of an oncological patient and healthy control, we identified the peptides that are more abundant in oncological patients than the healthy controls and vice versa. This way, we identified 40 spectra of peptides among which 20 spectra were most frequent in oncological samples and remaining 20 spectra were more abundant in healthy controls. In this study, we used these 40 peptide spectra as a feature set (called fset40) for developing models for predicting oncological status (Figure 2). As shown in Table 2, we achieved maximum performance using SVM-based models of SVM*^light^* (MCC of 0.77 with 88.31% accuracy) and SMO-Puk from Weka (MCC of 0.77 with 88.24% accuracy). This performance is lower than the performance we achieved using fset173; it may be owed to less number of features.

**Table 2:** The performance of models developed using 40 peptide spectra in fset40; models developed using various machine learning techniques.

| Method | Threshold | Sensitivity | Specificity | Accuracy | MCC |
| --- | --- | --- | --- | --- | --- |
| **Random-forest** | 0.5 | 89.57 | 80.24 | 84.94 | 0.70 |
| **SMO-Polykernel** | 0.5 | 87.15 | 85.23 | 86.20 | 0.72 |
| **SMO-Puk** | 0.5 | 88.85 | 85.03 | 86.96 | 0.74 |
| **SMO-RBF** | 0.5 | 86.56 | 85.23 | 85.90 | 0.72 |
| **SVM** | **0** | **90.03** | **86.56** | **88.31** | **0.77** |

**MCC: Matthews Correlation Coefficient**

**Figure 2:**
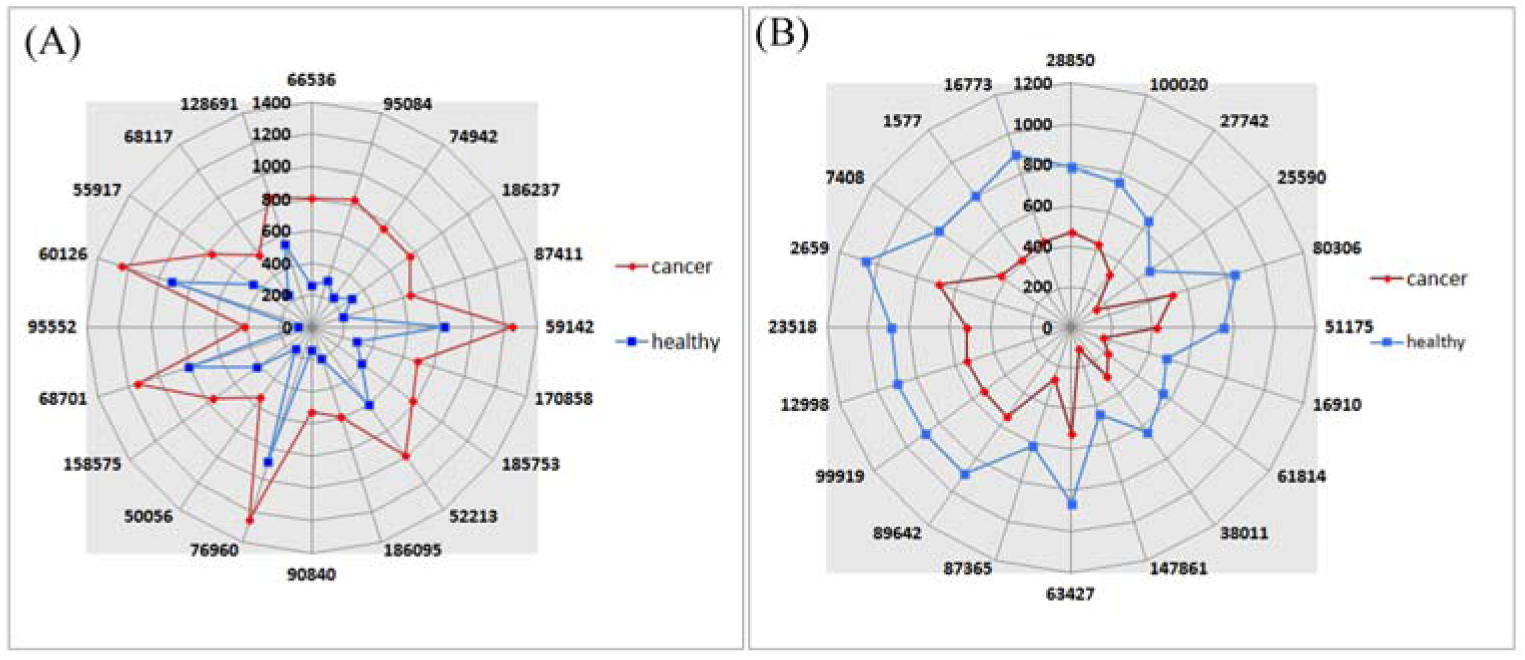
The radar graph shows the top peptide IDs that are (A) more frequent in oncological samples (red) as compared to healthy controls (blue) (B) more frequent in healthy controls (blue) as compared to oncological samples (red).

### 3.2 Peptide level prediction

The ideal biomarker panel is the one in which minimum number of peptides can differentiate well between oncological samples and healthy controls. In addition, the user is also interested to know the sequence of these peptides as spectra of peptide provide very limited information. Thus, we characterized 173 peptides spectra in fset173 and found that authors have annotated 61 peptides with their amino acid sequence and post-translational modifications. The propensities of these well-annotated 61 peptides were used for developing models for predicting oncological status.

The feature set of 61 peptides is referred to as fset61 in this study. As shown in Table 3, the performance of SVM*^light^* based model attained the accuracy of 87.75% with 0.76 MCC.

**Table 3:** The comparative performance of models developed using different machine learning techniques. The feature set fset61 contains spectra of 61 peptides used for building models.

| Method | Threshold | Sensitivity | Specificity | Accuracy | MCC |
| --- | --- | --- | --- | --- | --- |
| **Random-forest** | 0.5 | 89.38 | 78.58 | 84.02 | 0.68 |
| **SMO-Polykernel** | 0.5 | 85.38 | 84.63 | 85.01 | 0.70 |
| **SMO-Puk** | 0.5 | 88.33 | 84.10 | 86.23 | 0.73 |
| **SMO-RBF** | 0.5 | 86.23 | 84.23 | 85.24 | 0.70 |
| **SVM** | **0** | **90.30** | **85.16** | **87.75** | **0.76** |

MCC: Matthews Correlation Coefficient

### 3.3 Protein level prediction

The protein level analysis ensures even if some of the peptides of a protein are not detectable in the sample, average expression of all the peptide fragments belonging to a protein will well discriminate the samples. Therefore, we developed protein level biomarkers that can infer the oncological status of the patient. Out of 5605 peptides, we got detailed information about 953 peptides that includes their sequence, source protein and/or post-translational modifications. These peptides are derived from 118 proteins, out of which 69 proteins have two or more than two peptide fragments. In this study, feature set of 69 proteins is referred to as fset69. We developed SVM-based models using log-transformed expression/quantity of these proteins or oncological biomarkers. In order to compute quantity (or expression) of protein in a sample, we took mean of intensities of peptides (or fragment of proteins) belonging to that protein. The intensity of a peptide is proportional to the quantity of peptide in a sample. Similarly, we computed median and maximum expression of each protein (see Methods). The performance of SVM models developed using protein expression is displayed in Table 4. We achieved the highest accuracy of 85.27% with 0.71 MCC, where protein expression is represented by mean and maximum peptide intensities. These results indicate that quantity/expression of a protein in a sample can also be used for predicting oncological status.

**Table 4:** The performance of models developed using expression of 69 proteins, computed from intensities of their peptides. These models were developed on protein feature set, fset69.

| Protein expression | Threshold | Sensitivity | Specificity | Accuracy | MCC |
| --- | --- | --- | --- | --- | --- |
| **Mean** | 0 | 86.43 | 84.10 | 85.27 | 0.71 |
| **Median** | 0 | 82.75 | 81.84 | 82.30 | 0.65 |
| **Maximum** | 0 | 88.13 | 82.37 | 85.27 | 0.71 |

MCC: Matthews Correlation Coefficient

In order to reduce the number of proteins that can detect the oncological status of a urine sample, we identified proteins corresponding to a selected set of peptides. We obtained 61 sequences corresponding to peptides in fset173. We mapped these 61 peptide sequences on proteins; these peptides were mapped to 15 proteins such as CO1A1_HUMAN, CO3A1_HUMAN, CO1A2_HUMAN, COIA1_HUMAN, A1AT_HUMAN, UROM_HUMAN, HBB_HUMAN, *etc*. Among these 15 proteins, nine proteins have two or more than two peptides in the dataset. The model developed using the mean expression of 9 proteins (fset9) achieved maximum MCC of 0.63 with accuracy 81.27% (Table 5).

**Table 5:**
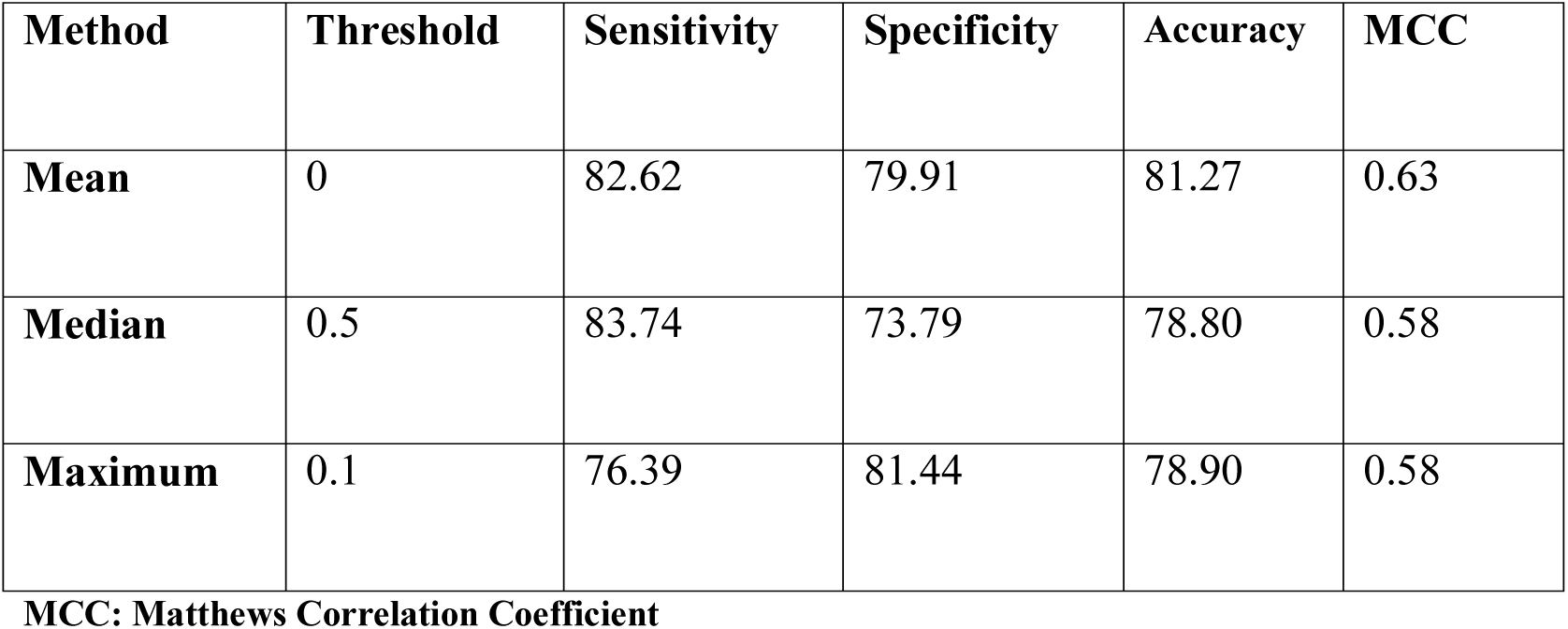
The performance of models developed using expression of 9 proteins, developed on feature set, fset9.

In order to infer the contribution of each protein, we developed a model for every protein using simple expression based cut-off using different thresholds. As shown in Table 6, we achieved maximum MCC of 0.57 with 78.17% accuracy for protein CO1A1_HUMAN, which accords with the previous observations that collagen fragments are potential candidate biomarkers for measuring oncological disorders.

**Table 6:**
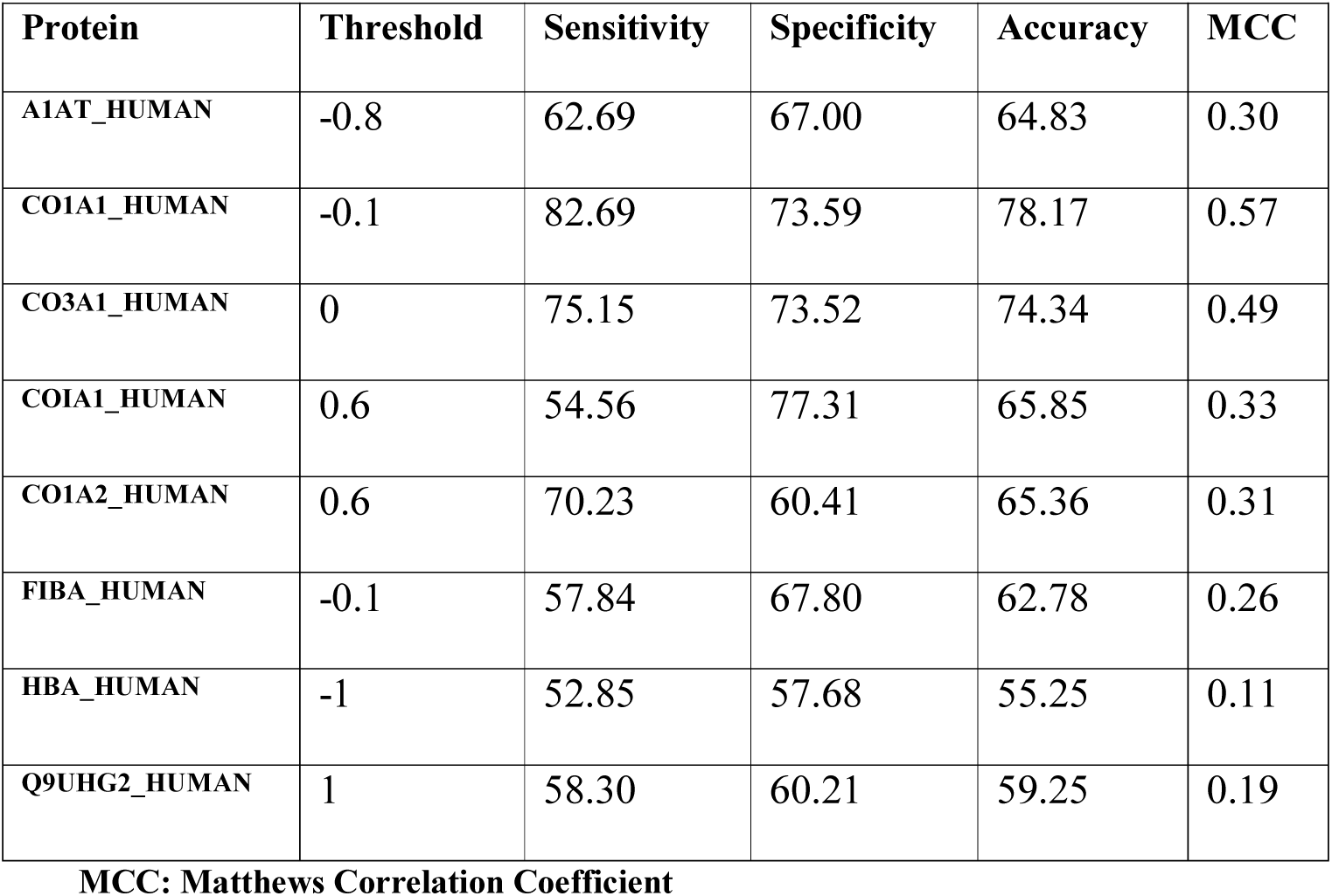
The performance of prediction methods based on expression of individual proteins. This is a simple threshold-based method developed for each protein in feature set, fset9.

## 4 Comparison to existing methods

It is important for any bioinformatics study to compare the newly developed method with the previously developed methods. Direct comparison of this method with previous studies is not possible because our models have been trained on the large number of urine samples to predict the oncological status. In 2009, Schiffer et al[16] developed model to predict the risk of muscle-invasive urothelial bladder carcinoma. Their model was trained on 424 urine samples (127 oncological patients and 297 healthy individuals) using panel of four peptides and achieved sensitivity of 81% with 57% specificity. In another study, Theodorescu et al[17] trained their model on 264 samples using 12 peptides biomarkers and achieved 91% sensitivity with 69% specificity. It is clear from above results that our models got comparable or better performance on large samples.

## 5 Web server implementation

In order to assist the scientific community, we developed a web server, CancerUBM (Urine-based BioMarkers for Oncological disorders) that implements models developed in this study for analysis and prediction of oncological status from proteomics data. This server has two major modules; one for predicting oncological status and another for analysis of proteomics data.

**Prediction Module**. This module allows the users to predict oncological status of their sample using three options; i) Mass-CE Spectra, ii) Peptide Sequence and iii) Protein Expression. In case of the first option, ‘Mass-CE Spectra’, the user needs to provide spectral details of each peptide that include its calibrated molecular mass, normalized CE migration time and intensity of spectra in a given sample. In case of ‘Peptide Sequence’ option, the user needs to provide calibrated molecular mass, normalized CE migration time, peptide sequence and intensity of peptide of the sample. Similarly in ‘Protein Expression’ module, server allows detecting oncological status of a sample from spectral intensities of peptides in a given protein. In summary, this module allows the user to predict oncological status of samples from proteomics data either based on spectra, peptide sequence or protein expression.

**Data Analysis Module**. This module allows the user to analyze their proteomics data at the protein and the peptide level. In case of the protein level option, server analyzes the proteomics data and computes mean, median and maximum intensity or expression of proteins along with graphical output. In case of peptide level analysis, server performs various analyses on peptide spectral intensities. For a particular peptide characterized by mass and CE time, this module gives the number of healthy and oncological samples observed in human urinary peptide database. It also provides the mean intensity of the peptide spectra across oncological samples and healthy controls. It also compares user’s data with the human urinary peptide database and gives the individual propensity of the peptide to contribute as the oncological biomarker and also gives the overall propensity to the user sample to be oncological. This web bench is available from URL, http://webs.iiitd.edu.in/raghava/cancerubm/ for public use.

## 6 Discussion

The utmost non-invasive and easy to attain body fluid for biomarker measurement is urine[20]. An inclusive understanding of the biomarkers is essential to timely predict the oncological disorders. Urine can be used to mine the important peptide biomarkers, which can predict the oncological status of the patient non-invasively. Human urinary peptide database represents a plethora of information that can be mined for detecting biomarkers for various diseases. In the present study, we have made a systematic effort to develop *in silico* models for the prediction of oncological state of the sample from urine proteomics data. Our primary analyses of the data indicate that nine proteins namely, A1AT_HUMAN, CO1A1_HUMAN, CO1A2_HUMAN, CO3A1_HUMAN, CO8A1_HUMAN, FIBA_HUMAN, HBB_HUMAN, Q9UHG2_HUMAN, UROM_HUMAN can act as potential biomarkers for detecting oncological sample of urine. Collagen fragments as oncological biomarkers have also been reported earlier in the literature. It has been shown that CO1A1_HUMAN has been linked with the bone metastasis[21] in breast cancer and is associated with angiogenesis and tumor growth[22]. In the past, elevated levels of FIBA_HUMAN chain has been shown to be associated with cancer progression[23]. These studies support the panel of biomarkers predicted for cancer in our analysis.

Our analysis, at peptide level, indicates that there are peptides that are more prominent in the urine sample of cancer patients. We used these peptides as biomarkers for developing models using various machine learning techniques. In our study, we achieved the best performance of models developed using SVM technique. We also developed models with biomarkers based on well-annotated peptides and source proteins. One of the limitations of this study is that details of clinical parameters of samples such as age, sex has not been taken into consideration in this study, as it is not given in source data. But as the sample number is quite large therefore this study can give the important insights of how urine can act as a good source for diagnostic peptides related to oncological disorders. In summary, we have made a systematic attempt to identify the potential biomarkers and developed highly accurate models for the prediction of oncological samples.

## 7 Concluding remarks

The *in silico* method, CancerUBM, can be useful for predicting urine peptide biomarkers, which can discriminate healthy controls from the oncological samples on the basis of the urine mass spectrometric data. The study also demonstrates that fragments of collagen can act as biomarkers to differentiate oncological samples and healthy controls. The cancer biomarkers identified (at three levels: spectra, peptide and protein) will be useful to understand why a particular set of proteins is more abundant in oncological patients. Our web bench will not only provide an oncological status of samples but also helps users in the identification of oncological biomarkers in their proteomics data. Thus, this open source web bench is expected to make a long-term impact in the field of oncological biology.

## 8 Acknowledgements

The authors are thankful to the Council of Scientific and Industrial Research (CSIR) and Indian Council of Medical Research (ICMR), Government of India, for financial assistance. The funders had no role in study design, data collection and analysis, decision to publish, or preparation of the manuscript.

## 9 Contributions

S.B. and K.C. compiled and organized the data. S.B. and K.C. developed the web interface and integrated the tools. S.B. and K.C. analyzed the data. S.B., K.C., A.G., G.P.S.R. wrote the manuscript. G.P.S.R. conceived the idea and coordinated the project. All coauthors have seen a draft copy of the manuscript and agree with its publication.

